# CHAMP delivers accurate taxonomic profiles of the prokaryotes, eukaryotes, and bacteriophages in the human microbiome

**DOI:** 10.1101/2024.08.15.608036

**Authors:** Sara Pita, Pernille Neve Myers, Joachim Johansen, Jakob Russel, Mads Cort Nielsen, Aron C. Eklund, Henrik Bjørn Nielsen

## Abstract

Accurate taxonomic profiling of the human microbiome composition is crucial for linking microbial species to health outcomes. Therefore, we created the Clinical Microbiomics Human Microbiome Profiler (CHAMP), a comprehensive tool designed for the profiling of prokaryotes, eukaryotes, and viruses across all body sites. Boasting a reference database derived from 30,382 human microbiome samples, CHAMP covers 6,567 prokaryotic and 244 eukaryotic species as well as 64,003 viruses. We used a diverse set of *in silico* metagenomes and DNA mock communities to benchmark CHAMP against the established profiling tools MetaPhlAn 4, Bracken 2, mOTUs 3, and Phanta. CHAMP demonstrated unparalleled species recall, F1 score, and significantly reduced false positives compared to all other tools benchmarked. Indeed, the false positive relative abundance (FPRA) for CHAMP was, on average, 50-fold lower than the second-best performing profiler. CHAMP also proved to be more robust than the other tools to low sequencing depths, highlighting its application for low biomass samples. Taken together, this establishes CHAMP as a best-in-class human microbiome profiler of prokaryotes, eukaryotes, and viruses in diverse and complex communities across low and high biomass samples. CHAMP profiling is offered as a service by Clinical Microbiomics A/S and available for a fee at https://cosmosidhub.com.

## Introduction

Understanding the role of the human microbiome in health and disease hinges on accurate species-level taxonomic profiling. This requires an accurate representation of the diverse consortium of bacteria, archaea, eukaryotes, and viruses across various human body sites (Huttenhower et al., 2012; Liang and Bushman, 2021). Hence, taxonomic profiling tools must accurately represent the true sample composition, even in samples with variable biomass and sequencing depths, while minimizing false detections.

The landscape of known human microbiome microorganisms has expanded significantly, largely due to the generation of bacterial and archaeal genome assemblies from metagenomic data, also known as metagenome-assembled genomes (MAGs). This has allowed the creation of comprehensive MAG databases such as the Unified Human Gastrointestinal Genome (UHGC) collection (Almeida et al., 2020) and the Early-Life Gut Genomes (ELGG) catalog (Zeng et al., 2022). Similarly, bacteriophage identification from metagenomic and metavirome data has led to the discovery of thousands of novel phages (Johansen et al., 2022). Binning of eukaryotic contigs from metagenomic samples, albeit less developed, has also yielded some MAGs, although nowhere near the number found for prokaryotes (Olm et al., 2019; Saraiva et al., 2023). Consequently, most profiling databases combine known and novel prokaryotic and viral species from isolates and metagenomes, whilst eukaryote genomes are primarily represented by sequenced isolates (Parks et al., 2021; Blanco-Míguez et al., 2023).

Multiple strategies for taxonomic profiling exist, among which the most used are marker-gene and k-mer based methods (Wright et al., 2023). Marker-gene based methods such as MetaPhlAn 4 (Blanco-Míguez et al., 2023) and MOTUs 3 (Ruscheweyh et al., 2022) align entire sequencing reads to specific gene catalogs, whereas k-mer based methods, such as Bracken 2, compare short subsequences of reads against extensive reference databases (Wood et al., 2019). Marker-gene and k-mer based strategies have both performed well in profiling benchmarks (Wright et al., 2023). Standardization of benchmarking practices in human microbiome research including convergence of the datasets (both *in silico* and DNA mock communities), performance metrics and reporting systems is crucial for the impartial evaluation of accuracy and precision across taxonomic profilers (Sczyrba et al., 2017; Amos et al., 2020).

Here we introduce CHAMP, a comprehensive human microbiome profiler, which synergizes sensitive marker gene identification with the potential of MAGs to profile prokaryotes and eukaryotes, and a specialized k-mer method for low-abundance phage detection. CHAMP was designed exclusively for the taxonomic profiling of short-read (>100bp), paired-end and human microbiome data. CHAMP focuses on accurately capturing the diversity across the human microbiome, covering a total of nine body sites and 6,546 bacterial species, 21 archaeal species, and 244 eukaryotic species, and 64,003 viruses. Our profiler shows improved performance across all domains of life and viruses, and is robust across sequencing depths, as validated by both *in silico* and sequenced DNA mock communities.

## Materials and Methods

### Human Microbiome Reference gene catalog for prokaryote and eukaryote profiling

The Clinical Microbiomics Human Microbiome Reference (HMR05) gene catalog was derived primarily from high-quality (HQ) prokaryotic MAGs identified *de novo.* These were complemented with MAGs from public repositories UHGC (82,834 MAGs and 6,428 isolates; Almeida et al., 2020) and ELGG (25,303 MAGs; Zeng et al., 2022). The *de novo* MAGs were generated from 30,382 human microbiome samples collected across nine distinct human body sites, including: gut (n = 19,296), small intestinal biopsies (n = 780), oral (n = 4,994), skin (n = 4,306), urine (n = 934), nasopharyngeal (n = 422), vaginal (n = 422), airway (n = 108), and milk (n = 100). 11% (n = 3,317) of the samples were not publicly available. In addition, isolate genome assemblies from NCBI (Sayers et al., 2022) and PATRIC (Wattam et al., 2017) were added to capture otherwise missing species of interest, including human-associated pathogens and probiotics. Human-relevant eukaryotic species were manually collected from various sources, including an analysis of gut fungal species (Nash et al., 2017), publicly available lists of pathogens from the World Health Organization (WHO), and the eukaryotes profiled by MetaPhlAn 4 (Blanco-Míguez et al., 2023), resulting in 2740 genomes representing 244 species. In addition, genomes from prokaryotic and eukaryotic species relevant for benchmarking and absolute abundance estimation by spike-in were also included.

For MAGs not obtained from publicly available MAG repositories, reads were trimmed, host-filtered, and assembled into contigs with MEGAHIT (v. 1.2.9, Li et al., 2015) or metaSPAdes (v. 3.15.5, Nurk et al., 2017), and then binned using VAMB (v. 3.0.6, Nissen et al., 2021). MAGs were considered high-quality if they had > 90% completeness and < 5% contamination based on CheckM2 (v. 2022-07-19; Chklovski et al., 2023) and passed the GUNC chimerism test (v. 1.0.5, Orakov et al., 2021).

Redundant MAGs were removed by clustering MAGs within a species cluster using a 99.5% ANI cutoff. All MAGs were taxonomically annotated using GTDB-Tk (v. 2.3.0, Chaumeil et al., 2022) with Genome Taxonomy Database (GTDB) release 214 (Parks et al., 2022). To combine MAGs from multiple VAMB batches and MAG repositories, MAGs annotated to the same species were merged. MAGs without GTDB-Tk species-level annotations were clustered into species clusters based on 95% average nucleotide identity (ANI) (dRep, Olm et al., 2017; FastANI, Jain et al., 2018). This resulted in 6,567 prokaryotic species clusters, 11% of which were unannotated at a species level according to GTDB.

### Identification of pan-genomes and signature genes

To derive a pan-genome catalog for each species, we called genes using Pyrodigal (v 2.0.2, Larralde, 2022) and then used a three-step clustering approach. First, nucleotide sequences of the genes were clustered with MMseqs2 (v. 14, Steinegger & Söding, 2018) with 98% identity and 90% bi-directional coverage. Second, the representatives from the first iteration were clustered with MMseqs2 to 95% identity and 90% bi-directional coverage. Representatives of the second iteration were chosen as the ones with highest cardinality from the first iteration. Third, the second iteration representatives were clustered with CD-HIT (cd-hit-est, v. 4.8.1, Fu et al., 2012) to 95% identity and 90% coverage of the shorter sequence. For the third iteration clusters, genes were discarded if they were overlapping with >20% of their length with another gene of higher cardinality, such that only non-overlapping (<20% of shorter sequence) genes with the highest cardinality were left. Genes shorter than 100 bp or with a species prevalence < 1% were discarded.

For prokaryotes and eukaryotes separately, the complete set of pangenomes were then clustered with MMseqs2 to 97% identity and 90% bi-directional coverage to obtain between-species clusters. The pan-genomes from prokaryotic (n = 6,567) and eukaryotic (n = 244) species were merged into a final gene catalog of 25,761,278 genes.

To enable quantification of each species in the database, up to 250 signature (also called marker) genes were selected for each species based on core genes (≥ 60% prevalence in species MAGs) with a length ≥ 200 bp and ≤ 20 kbp. To ensure specificity, a potential signature gene was removed if it aligned to another gene in the catalog with > 97% identity over 100 bp. However, if fewer than 20 genes met these criteria for a species, then genes with segments > 200 bp without alignments to other genes were used, and non-unique segments of these genes were masked.

### Preprocessing of sequencing reads

Raw reads were trimmed to remove adapters and low-quality bases (Phred score < 30) using AdapterRemoval (v. 2.3.1, Schubert et al., 2016). Then, host contamination was removed by discarding read pairs where either read mapped to the human reference genome GRCh38 with Bowtie2 (v. 2.4.2, Langmead & Salzberg, 2012). Resulting read pairs were retained if both reads had a length of at least 100 bp; these were classified as high-quality non-host (HQNH) reads.

### Abundance profiling of prokaryotes and eukaryotes

HQNH reads were mapped to the HMR05 pan-genome catalog using BWA mem (v. 0.7.17, Li and Durbin, 2009). An individual read was considered uniquely mapped to a gene if the MAPQ was ≥ 20 and the read aligned with ≥ 95 % identity over ≥ 100 bp.

However, if > 10 bases of the read did not align to the gene or extend beyond the gene, the read was considered unmapped. The expected read counts for signature genes in each species in each sample were modelled with a negative binomial distribution as follows. First, if (1) ≥50 of the signature genes for a species had non-zero read counts and (2) ≥99% of genes were expected to have non-zero read counts given the total read count for that species (1-(((n_genes-1)/n_genes)^n_reads) ≥0.99), then signature genes with zero reads were ignored in that sample. Second, the expected 99% quantile (between 0.5% and 99.5%) of read counts was calculated for each gene based on a negative binomial distribution with a mean proportional to the effective gene length and dispersion defined as log2 (effective gene length). The abundance of each species was then calculated, using only the signature genes with observed read counts within the expected 99% quantile, as the mean read count normalized by effective gene length. Species abundances were set to zero if fewer than 5 genes with non-zero read counts were within the 99% quantile. Furthermore, species with < 66% of genes with non-zero read counts within the 99% quantile were set to zero, unless the median abundance of signature genes was non-zero, in which case the median gene-length-corrected abundance of non-zero genes was used. Abundances were then normalized sample-wise such that the total abundance of all species sums to 100%.

### Human phage and viral genome database for bacteriophage profiling

The Human Virome Reference v. 1 (HVR01) was constructed by combining viral genomes identified *de novo* with the public viral databases: MGV (n = 54,118, Nayfach, Páez-Espino, et al., 2021), IMG/VR (n = 250,970, v. 4, downloaded 2022-12-19, using the subset of human-associated virus genomes, a database of infant viruses from the COPSAC consortium (n = 10,021, (Shah et al., 2023), and NCBI RefSeq viruses (n=15,247, downloaded 2023-03-02, Brister et al., 2015). Putative viral contigs from HMR05 MAGs were determined by running geNomad (v. 1.3.3, Camargo & Kyrpides, 2023) and accepting only hits with a geNomad viral score of ≥ 0.7. Putative viral contigs were retained if they had either (1) direct-terminal-repeats (DTR) and a minimum size of 2,000 bp, or (2) at least one viral hallmark gene and a minimum size of 4,000 bp. For datasets processed using VAMB metagenomic binning, multiple viral contigs were combined into a viral MAG if found in the same VAMB bin. Altogether, *de novo* discovery of viruses resulted in 547,573 putative viral genomes.

Each of the putative viral genomes were scored with a viral confidence score as described in the IMG/VR v4 (Camargo et al., 2023). First, viral genomes were annotated with CheckV (v.1.0.1, default settings, Nayfach et al., 2021a). Second, viral genomes were clustered with RefSeq viral genomes into viral operational taxonomic units (vOTU) based on all-vs-all genome alignment using Blastn (v 2.10.1+, -task megablast -evalue 1e-5 -max_target_seqs 10000, McGinnis and Madden, 2004).

Genomes were clustered into a vOTU if they had a 95% ANI across at least 85% of the genome. Subsequently, viral genomes were scored accordingly:

3 points: Viral genome clustered with a RefSeq viral genome into a vOTU, and three or more geNomad viral hallmark markers.

2 points: High-confidence CheckV average amino acid identity (AAI) completeness estimate, and at least two geNomad viral hallmark markers.

1 point: Medium-confidence CheckV AAI completeness estimate, and one geNomad viral hallmark marker; direct or inverted terminal repeats; two or more matches to CRISPR spacers.

Viral genomes with at least 2 points (n = 231,587 genomes) were retained for downstream analysis. These genomes were combined with viral genomes from the public viral databases and clustered into vOTUs (95% ANI across 85% aligned genome fraction [AF]) to form a non-redundant viral genome database. To optimize the non-redundant database for viral profiling, we minimized the degree of shared nucleotide sequences between representatives by filtering out vOTUs as follows: (1) viral genomes in a vOTU (n > 1) with any edges to genomes of other vOTUs were removed from that cluster; (2) satellite vOTUs (n = 1, one genome in cluster) with edges to multiple vOTUs were removed entirely; (3) satellite vOTUs (n = 1) with edges to members of only a single other vOTU were reassigned to that vOTU; (4) entire vOTUs were removed if genome members had borderline similarity to other clusters (ANI > 93 and 30 < AF < 85) or (93 < ANI < 95 and AF > 85 AF). The longest genome within each of the refined vOTUs was chosen as the representative genome, resulting in a final representative viral database containing 64,003 genomes.

Genomes acquired from RefSeq and the COPSAC consortium were assigned their preexisting taxonomy. The remaining uncultivated viral genomes without taxonomic annotation were assigned to viral taxa as defined in the International Committee on Taxonomy of Viruses (ICTV) Release #38 (Nayfach et al., 2021b). The following taxonomic classification methods were used in order of priority: (1) clustering with RefSeq viral genomes; (2) clustering with COPSAC viral genomes; (3) geNomad marker-based taxonomic assignment. For the marker-based assignment, geNomad was employed to classify sequences using taxonomically informative protein profiles. The first taxonomic annotation steps (1-3) yielded class and family annotation for 94.7% and 12.8% of viruses, respectively, of which 49,997 out of 64,003 genomes belonged to the class Caudoviricetes, *i.e.,* tailed bacteriophages.

### Prediction of vOTU host taxonomy

The bacterial and archaeal-host taxonomy for each vOTU was predicted at each taxonomic level based on CRISPR spacer alignment and k-mer matching between vOTUs and prokaryotic genomes.

For the CRISPR spacers, a non-redundant database (100% identity and 100% coverage) of 4,793,298 CRISPR spacers, originally derived from approximately 1.6 million genomes from the NCBI and MAG study (Camargo et al., 2023) were merged with a non-redundant database (100% identity and 100% coverage) of 1,049,986 CRISPR spacers extracted using minced (v 0.4.2, Bland et al., 2007) from the HMR05 HQ MAGs. The spacers of the combined database were mapped with blastn (v.2.10.1+, - max_target_seqs = 1000 -word_size = 8 -dust = no) against all representative vOTU genomes. Confident virus-host connections were considered if spacer-alignments had a length of at least 25 bp and 95% spacer coverage with a maximum of 1 mismatch.

K-mer matching was performed using PHIST (v. 1.0.0, default settings, Zielezinski et al., 2022) between representative vOTU genomes and bacterial and archaeal MAGs from HMR05 HQ MAGs. To prevent spurious matches, only virus-host connections where at least 20% of the viral k-mers were found in a prokaryotic genome were accepted.

The prokaryotic host taxon was then assigned to each vOTU at the lowest taxonomic rank, having at least two connections (either spacers or k-mers) and representing > 50% of all connections. We chose the method that was supported by the greatest number of host assignments, prioritizing CRISPR-spacers over k-mer matching annotations.

### Abundance profiling of bacteriophages

Read pairs mapping to each vOTU genome were counted using KMA (v. 1.4.7, Clausen et al., 2018) with the settings: “-mrs 0.01 -apm p”. A vOTU was considered detected if it had ≥ 5 aligned paired-end reads and either (1) the percentage of identical nucleotides between genome and consensus sequence assembled from mapped reads was ≥ 99%, and ≥ 5% of the genome was covered by reads, or (2) the percentage of identical nucleotides between genome and consensus sequence of mapped reads was ≥ 95%, and ≥ 30% of the genome was covered by reads. vOTU abundance was estimated based on number of mapped read pairs divided by genome length.

### Benchmarking of prokaryotic profiling

For prokaryotic benchmarking, ten body site-representative prokaryotic metagenomes were simulated for each of the following five body sites: adult gut, infant gut, oral, skin, and vagina. Genome accession ids for prokaryotic species found in each human body site were identified from published literature to avoid biasing the benchmark to our own database (Bäckhed et al., 2015; Proctor et al., 2019; Saheb Kashaf et al., 2021). For the adult gut, oral and vagina, GCA/GCF identifiers were extracted from the publications and used for simulation if present in GTDB metadata. For the skin and infant gut, MAGs from the publications were classified with GTDB-Tk and the GCA/GCF references of classified MAGs were used for simulation (Supplementary Table 1). 100 (80 for vaginal) randomly sampled prokaryotic species specific to each body site were provided for each community. Metagenomes were generated using CAMISIM (Fritz et al., 2019), which simulates Gb of Illumina 2 ×150 bp paired end reads with the default HiSeq 2500 error profile and a mean insert size of 200 bp. To assess profiling performance for a range of sequencing depths, the 50 *in silico* metagenomes were also downsampled with seqtk (-s100) to sequencing depths of 20, 5, 2, 1, 0.5, 0.25 and 0.1 million read pairs.

Prokaryotic profiling was also benchmarked with DNA reference reagents. NIBSC sequencing data for ten DNA mock communities comprising 19 common gut bacterial species was downloaded from the NCBI Sequence Read Archive (NCBI Bioproject ID PRJNA622674, Amos et al., 2020).

CHAMP performance was compared against popular and third-party benchmarked top-ranking (Meyer et al., 2022; Poussin et al., 2022) open-source taxonomic profilers: marker-gene based profilers MetaPhlAn 4 (v 4.0.6, database from 2022-12, Blanco-Míguez et al., 2023) and mOTUs 3 (Ruscheweyh et al., 2022), and Bracken 2 (Lu et al., 2017), which uses the k-mer based method KRAKEN for mapping (Wood et al., 2019). Two iterations of Bracken 2 profiling were conducted: the first, using the standard Bracken 2 database based on the RefSeq database (O’Leary et al., 2016) and a second instance, using the GTDB release 214 (r214).

MetaPhlAn 4, mOTUs 3 and Bracken 2 were run on default settings. A long tail of incorrectly predicted low abundance species leading to low precision estimates has been previously shown for kmer-based profilers (Sun et al., 2021). To assess the veracity of low abundance species predictions, taxonomic profiles for the 50 *in silico* human body site metagenomes were filtered to relative abundance cutoffs of 0.0005, 0.0001, 0.0005, 0.001, 0.005 and 0.01 (Supplementary Figure 1) across profilers. Based on these results, we chose to filter relative abundances < 0.001 from Bracken 2 profiling output to maximize F1 scores for the prokaryotic benchmark. We did not filter low abundant species detections from any of the other profilers.

We also tested Bracken 2 built with GTDB r214 database to harmonize species annotations between Bracken 2 and CHAMP, however, Bracken 2 performed better on RefSeq than GTDB (Supplementary Figure 2). Hence, all other Bracken 2 benchmarks use the RefSeq database.

### Benchmarking of eukaryotic profiling

30 eukaryotic *in silico* metagenomes comprising up to 200 randomly sampled genomes were generated using CAMISIM. Species genomes were selected from a set of 113 eukaryotic species (Supplementary Table 2) corresponding to the eukaryotic species within both CHAMP (244 species) and MetaPhlAn 4 (122 species) databases. Eukaryotic taxonomy was based on NCBI annotations (Schoch et al., 2020). CHAMP eukaryotic profiling was compared against MetaPhlAn 4 using these 30 *in silico* eukaryotic metagenomes. mOTUs 3 and Bracken 2 were excluded from the eukaryotic benchmark, as their default databases do not include eukaryotes.

### Benchmarking of phage profiling using i*n silico* mixed prokaryotic and phage communities

The phage-inclusive profiling of CHAMP was benchmarked against Phanta (Pinto et al., 2023) using 10 synthetic metagenomic mixed community datasets constructed using CAMISIM (Fritz et al., 2019). Each community consisted of 200 randomly selected bacterial genomes from GTDB with species-level annotation and 200 viral genomes from the Gut Phage Database (GPD, Camarillo-Guerrero et al., 2021). GPD genomes were sourced to avoid simulating reads from genomes corresponding to genomes present in HVR01 or Phanta (matched using genome-index). GPD genomes were clustered with genomes in MGV vOTUs using ANI ≥ 95% and coverage ≥ 85% to identify the subset of genomes covered by both databases (MGV vOTUs were part of both HVR01 and Phanta). One GPD genome was sourced for each vOTU (Supplementary Table 3). ART Illumina HiSeq 2500 error profile with an insert size of 200 ± 25 bp was used to simulate a total of 50 million paired reads (2 × 150 bp) for each community with 95% of the reads originating from bacteria and 5% of the reads originating from viruses. Reads were subsampled for each community using seqtk (parameters: *-s100*) to generate metagenomes at 25, 20, and 10 million reads for each of the 10 communities, thereby bringing the total number of metagenomes in the benchmarking dataset to 40.

To investigate the robustness of CHAMP to lower sequencing depths, the 10 metagenomes were rarefied to 5, 2, 1, 0.5, 0.25, and 0.1 million reads.

### Benchmarking of phage profiling and removal of false-positives proviruses

Phanta (v.0.3.0) was run using default settings but changing bacterial and viral coverage threshold according to the sequencing depth of the profiled metagenome. The threshold between bacterial and viral coverage was set to 0.05/0.35 for *in-silico* samples with a depth of 50 million reads, 0.02/0.2 for samples with a depth between 20-25 million reads, and 0.01/0.05 for samples with a depth equal to 10 million reads.

*In silico* read simulations from whole bacterial genomes can introduce the presence of “false-positive” (FP) viruses, due to reads from prophages integrated in bacterial chromosomes mapping to viruses in the profiling database. As these prophages are not part of the expected true set of viruses in each benchmark community, we introduced a filtering criterion to detect these false positives (although they are true prophages) to allow a fair benchmark between profilers, in line with the methodology applied previously (Pinto et al., 2023). Viruses detected by either CHAMP or Phanta which attracted ten or more reads from a bacterial chromosome were classified as FP viruses and removed from the set of viruses detected in each sample.

### Classification performance metrics

Binary performance metrics were calculated using correctly (TP) and incorrectly (FP) detected species by a profiler along with the number of undetected species present in each sample (FN). Precision = TP/(TP+FP), describes the fraction of species detected by a classifier correctly, while recall = TP/(TP+FN), is the fraction of correctly detected species within a sample. The F1 score is the harmonic mean of recall and precision and was calculated as F1 = (2 × precision × recall)/(precision + recall). We also report false positive relative abundance (FPRA) = FP abundance/total species abundance; sensitivity = TP/(TP+FN); and Bray–Curtis similarity = 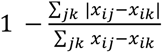, where *x* and *x* refer to the observed and ground truth abundance of a species as defined by NIBSC (Amos et al., 2020). We also calculated additional benchmarking metrics as described in (Meyer et al., 2019; Supplementary Table 4).

## Results

### Development of the Human Microbiome Reference and the Human Virome Reference databases

Large-scale efforts to build human microbiome references have predominantly focused on cataloging prokaryotic genomes from human stool samples (Almeida et al., 2019; Nayfach et al., 2019), with lesser emphasis on other body sites (Pasolli et al., 2019) and even less on identifying viral genomes (Benler et al., 2021; Shah et al., 2023). To broaden the spectrum of species and viral genomes represented, we compiled 4,994 oral, 4,306 skin, 422 vaginal, 934 urine, 422 nasopharyngeal, 108 airways, and 100 human milk samples. Notably, 30% of these samples were sourced from non-public domains. Furthermore, we included 780 samples from intestinal biopsies to capture the distinct microbiome of the gastrointestinal tract compared to stool (Zmora et al., 2018), 2,158 stool samples from countries unrepresented in the public repositories, and 1,000 stool samples from the SCAPIS cohort (Dekkers et al., 2022). These samples were batch processed using VAMB (Nissen et al.,2021) to bin contigs, resulting in 147,427 MAGs from prokaryotic species of which 79,796 were high-quality (HQ) and non-chimeric, along with, 547,573 putative viral genomes (see Materials and Methods).

We supplemented these 79,796 HQ, non-chimeric prokaryotic MAGs with MAGs from public repositories—UHGC (82,834 MAGs and 6,428 isolates, Almeida et al., 2020) and ELGG (25,303 MAGs, Zeng et al., 2022)—and with 27,492 genome assemblies from NCBI (Sayers et al., 2022) and PATRIC. This inclusion aimed to encompass missing species of interest such as human-associated pathogens, gut fungal species, probiotics, and species found in DNA mock communities or utilized as spike-ins for absolute abundance estimation. All prokaryotic MAGs and reference genomes were organized into species clusters based on GTDB annotations. MAGs without species-level annotation were clustered with a 95% ANI threshold to each other and to the representative MAGs with a species level annotation. Redundant MAGs within each cluster were pruned using a 99.5% ANI cutoff yielding a total of 102,445 prokaryotic MAGs. The 547,573 putative viral genomes from our human metagenomic data were merged with 330,356 viral genomes from four public viral repositories, clustered with a 95% ANI threshold and scored based on presence of viral features to form a comprehensive, non-redundant database with 64,003 viral genomes.

Altogether, the Human Microbiome Reference (HMR05) and the Human Virome Reference version 1 (HVR01) encompass 6,546 bacterial species, 21 archaeal species, 244 eukaryotic species, and 64,003 viruses.

### Benchmarking of prokaryotic profiling using CAMI *in silico* datasets

CHAMP employs signature genes for profiling prokaryotic and eukaryotic species. These are selected genes that are ubiquitous among strains of the species while not similar to genes of other species.

Nevertheless, there may exist rare genetic elements that result in irregular mapping to signature genes. For example, from rare species or rare conspecific genetic diversity. To counteract this, CHAMP models the abundance of each signature gene in every sample using a negative binominal distribution and subsequently filter out genes that fall outside the 99% percentile expected mapping, assuming that a given set of signature genes have the same abundance in a sample. The performance of CHAMP was evaluated by benchmarking it against three prominent metagenomic profilers: MetaPhlAn 4, mOTUs 3 and Bracken 2, across a variety of sample types using standardized benchmarking metrics (Meyer et al., 2019).

We first assessed the performance of CHAMP on simulated mock communities, designed to mirror the diversity of prokaryotic species within the human microbiome. These communities, representing five key human body sites: the adult gut, infant gut, oral cavity, skin, and vagina, and were generated using the CAMISIM tool (Fritz et al., 2019). We simulated 10 communities for each of the five body sites.

CHAMP excelled over all other profilers in recall (percentage of actual species that were detected correctly), across all body sites, indicating its superior ability to detect species. Both CHAMP and MetaPhlAn 4 demonstrated exceptional precision (the percentage of correctly detected species) across these community types, with average precisions of 0.95 ± 0.03 and 0.93 ± 0.03, respectively, significantly surpassing Bracken 2 and mOTUs 3 (Figure 1). CHAMP showed superior precision in the adult and infant gut, as well as oral communities, while MetaPhlAn 4 performed marginally better in skin and vaginal communities. Importantly, CHAMP struck the best precision-recall tradeoff with the highest F1 scores across all body sites. For similarity estimates (Bray–Curtis similarity a measure of abundance differences from the ground truth), CHAMP and MetaPhlAn 4 again achieved the highest averages (avg. 0.9 ± 0.08 vs. avg. 0.9 ± 0.08) across body site communities. Finally, CHAMP excelled with an FPRA (avg. 0.008 ± 0.02) that is more than six times less than the second-best profiler MetaPhlAn 4 (0.05 ± 0.08). Overall, CHAMP performed exceptionally well, outperforming all other profilers across key metrics and body sites. We found similar results in benchmarks at genus-level (Supplementary Figure 3).

**Figure 1.**
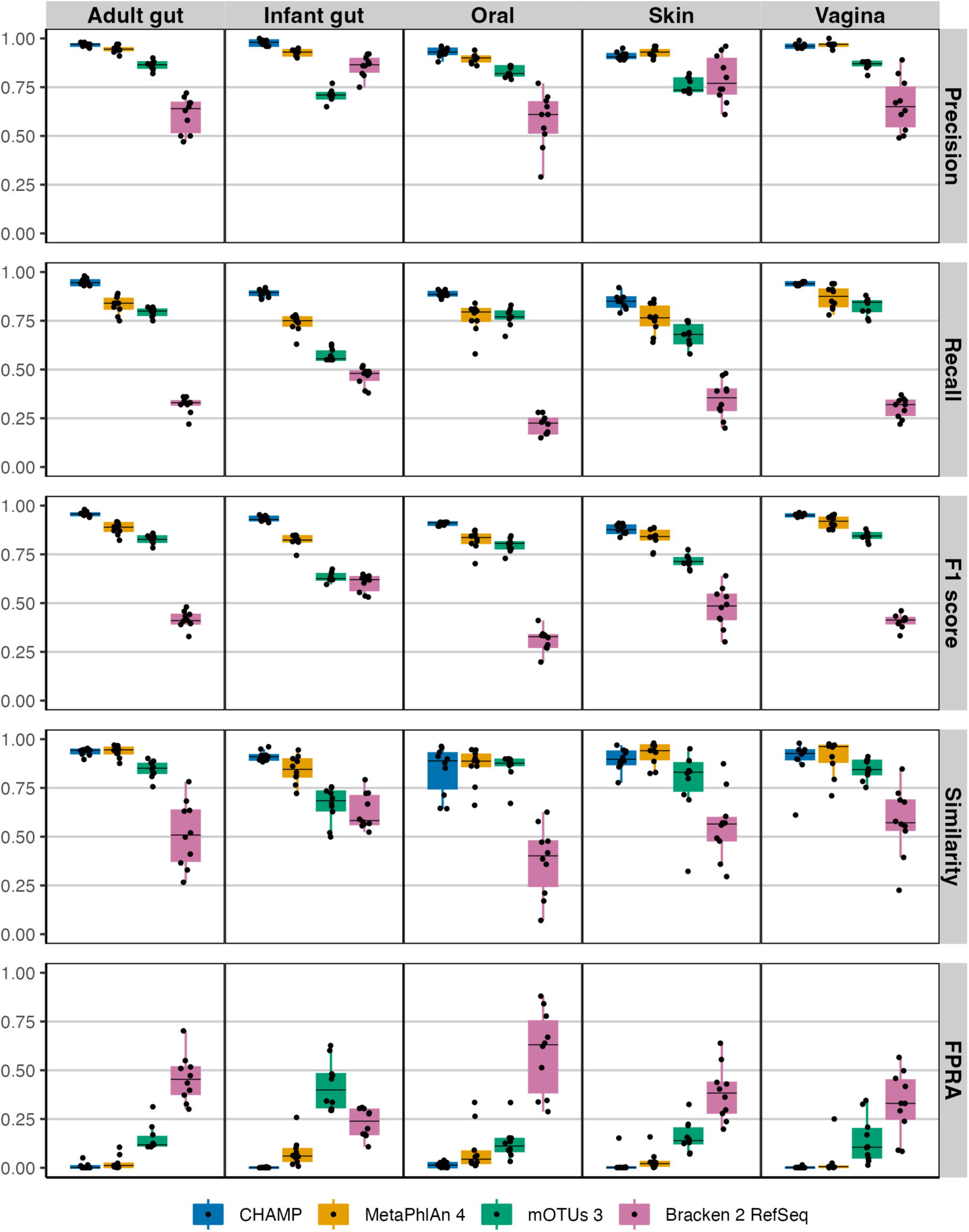
Evaluating the taxonomic profilers CHAMP, MetaPhlAn 4, mOTUs 3, and Bracken 2 on key benchmarking metrics: precision, recall, F1 score, similarity, and FPRA across five human body communities. A total of 50 metagenomes (10 per body site) each with 100 prokaryotic species were simulated using CAMISIM.

Given the significant variability in sequencing depth across studies and its known impact on taxonomic profiling, we evaluated the performance of CHAMP compared to the other profilers by generating *in-silico* metagenomes at seven different sequencing depths, ranging from 20 million to 100,000 reads (Figure 2). Precision was found to be consistent across all sequencing depths for each profiler. However, recall and, to some extent, F1 scores, declined with reduced sequencing depths.

**Figure 2.**
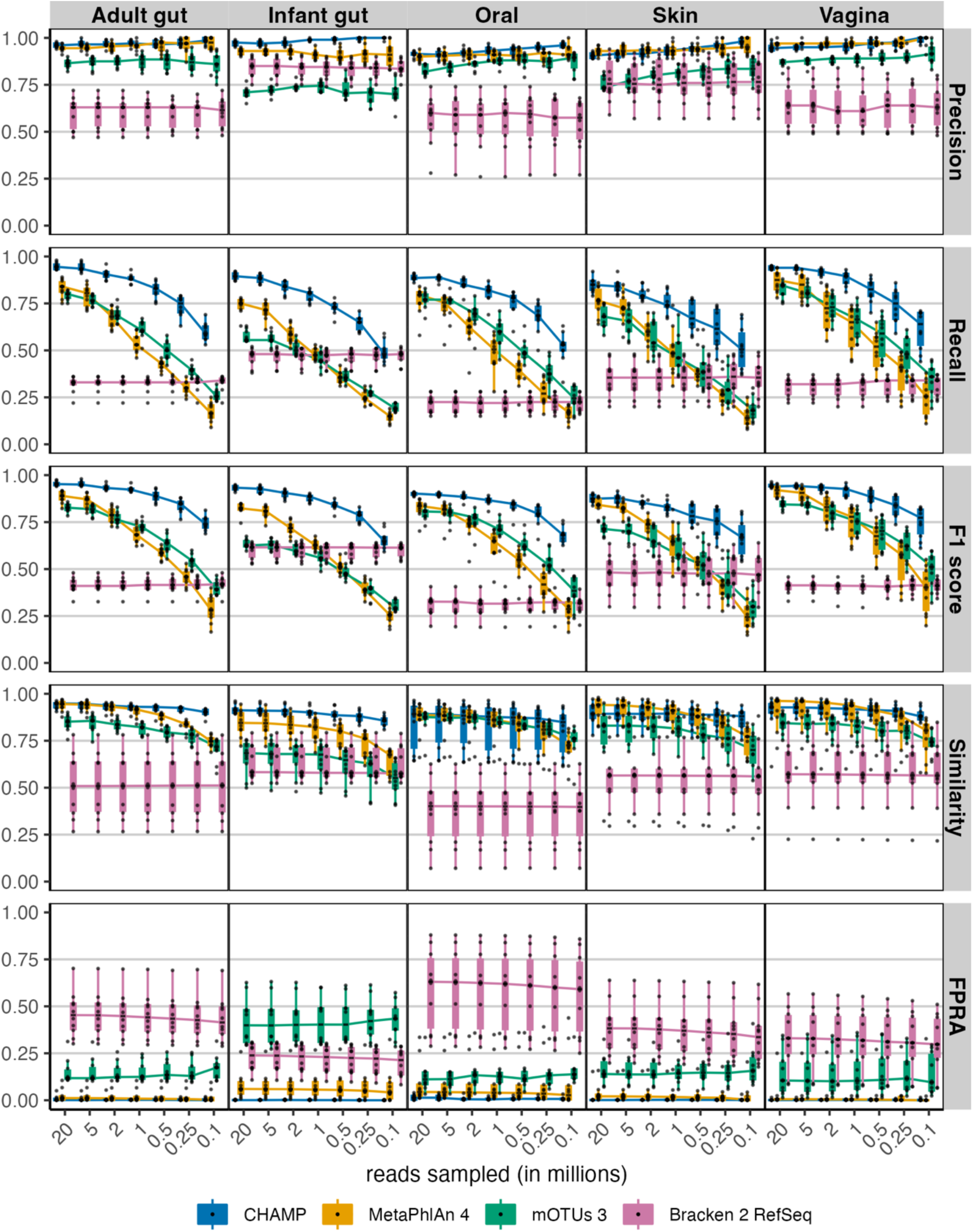
Evaluating performance across the sampled sequencing depths: 20, 5, 2, 1, 0.5, 0.25 and 0.1 million reads for the four taxonomic profilers: CHAMP, MetaPhlAn 4, mOTUs 3, and Bracken 2. The following benchmarking metrics were compared: precision, recall, F1 score, similarity, and FPRA. Each of 50 (10 per body site) metagenomes comprising 100 species were randomly sampled at seven sequencing depths resulting in 350 metagenomic samples.

Notably, CHAMP exhibited enhanced robustness at lower sequencing depths, particularly between 1 million and 100,000 paired reads. At the threshold of 500,000 paired reads, CHAMP had an average F1 score 1.5 times higher than that of the next-best profiler for the F1 score, MetaPhlAn 4, across all body site communities. This supports the utility of shallow shotgun sequencing as a cost-effective yet accurate approach for taxonomic profiling, where CHAMP at 500,000 reads, on average, had a 17-fold lower FPRA compared to the second-best profiler.

### Benchmarking prokaryotic profiling using NIBSC reference reagents

We assessed the performance of CHAMP against MetaPhlAn 4, Bracken 2, and mOTUs 3 on two DNA reference reagents provided by NIBSC: Gut-Hi-Low-RR, featuring even compositions, and Gut-Mix-RR, with staggered compositions of 19 common gut microbial species. Each reagent included five samples. For both DNA reference reagents, CHAMP showed the lowest average FPRA (Table 1), significantly outperforming the next-best profiler in terms of FPRA, by 55 times in Gut-Mix-RR and 110 times in Gut-Hi-Low-RR. CHAMP demonstrated superior sensitivity compared to MetaPhlAn 4 and mOTUs 3 on both reagents and outperformed Bracken 2 on Gut-Hi-Low-RR. For Gut-Mix-RR, Bracken 2 achieved perfect sensitivity, although it consistently detected more species than those that were in the reagents. Overall, CHAMP excelled in benchmarking metrics and showed unparalleled FPRA performance against current state-of-the-art profilers.

**Table 1.** Evaluation of CHAMP against other profilers using the NIBSC DNA Reference Reagents Gut-Hi-Low-RR and Gut-Mix-RR.

**A.**
| **<u>Gut-Hi-Low-RR</u>** |  |  |  |  |
| --- | --- | --- | --- | --- |
| **Profiler** | **Sensitivity** | **FPRA** | **Diversity** | **Similarity** |
| CHAMP | 94 | 0.01 | 19 | 0.62 |
| Bracken 2 | 91.6 | 4.26 | 21.4 | 0.76 |
| MetaPhlAn 4 | 89 | 1.1 | 18.8 | 0.65 |
| mOTUs 3 | 70 | 18.1 | 20.6 | 0.53 |

**B.**
| **<u>Gut-Mix-RR</u>** |  |  |  |  |
| --- | --- | --- | --- | --- |
| **Profiler** | **Sensitivity** | **FPRA** | **Diversity** | **Similarity** |
| Bracken 2 | 100 | 4.96 | 22.2 | 0.76 |
| CHAMP | 96 | 0.09 | 19.8 | 0.72 |
| MetaPhlAn 4 | 90 | 5.55 | 19 | 0.72 |
| mOTUs 3 | 70 | 26.2 | 24.2 | 0.63 |

### Benchmarking of eukaryotic profiling using CAMI *in silico* datasets

Eukaryotes, integral components of the human microbiome, have only recently been incorporated into many microbiome profilers (Blanco-Míguez et al., 2023). In assessing eukaryotic species-level profiling, we compared CHAMP with MetaPhlAn 4, excluding mOTUs 3 and Bracken 2 due to their lack of eukaryotic coverage. Using 113 eukaryotic species common between CHAMP and MetaPhlAn 4 databases, we generated 30 simulated mock communities for benchmarking. Both CHAMP and MetaPhlAn 4 exhibited high precision on these communities (avg. 0.97 ± 0.02 and 0.95 ± 0.03, respectively, Figure 3). CHAMP demonstrated superior recall compared to MetaPhlAn 4 (avg. 0.90 ± 0.05 vs. 0.53 ± 0.08), resulting in a higher F1 score average of 0.93 ± 0.03 compared to 0.67 ± 0.07 for MetaPhlAn 4. Additionally, CHAMP surpassed MetaPhlAn 4 in the similarity of predicted profiles to the ground truth composition and had lower abundance of false positives (FPRA avg. 0.02 ± 0.06 vs. 0.03 ± 0.03). Altogether, CHAMP displayed superior performance at eukaryotic profiling compared to MetaPhlAn 4 across all benchmarking metrics.

**Figure 3.**
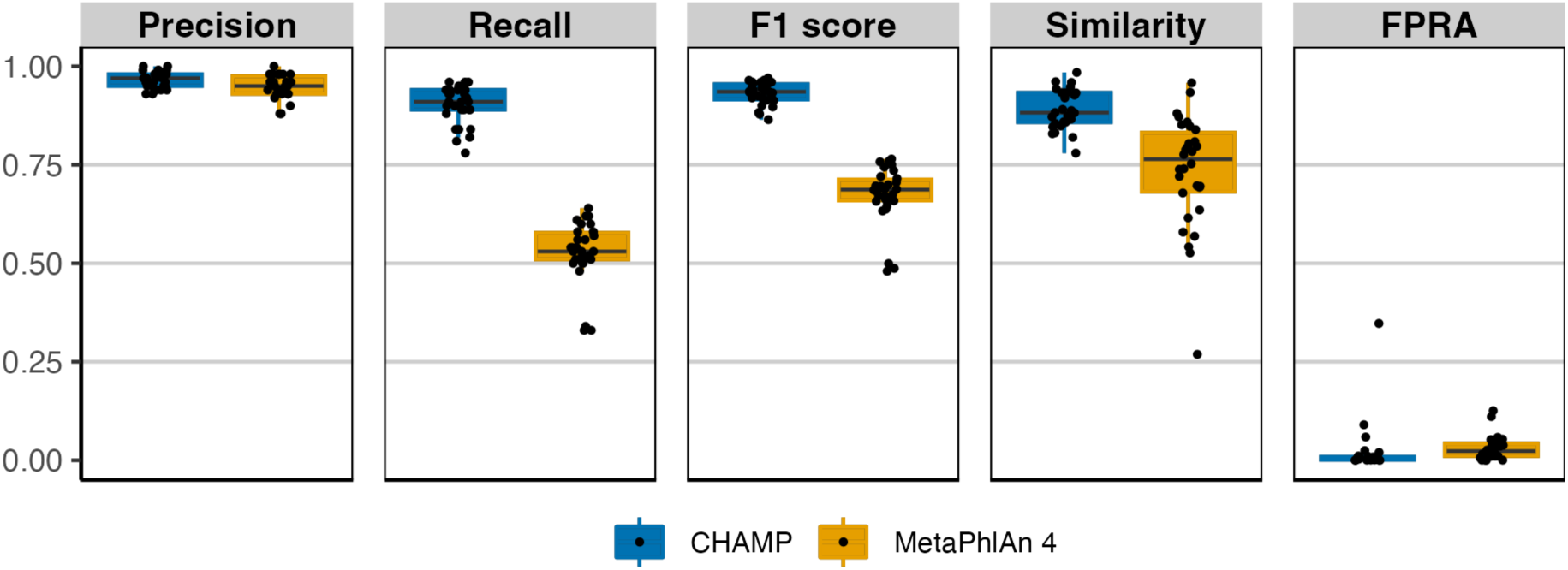
Comparison of eukaryotic taxonomic profiling by CHAMP and MetaPhlAn 4 using *in silico* metagenomes. 30 metagenomes were simulated encompassing 113 eukaryotic species found both in CHAMP and MetaPhlAn 4 databases. Benchmarking metrics included: precision, recall, F1 score, similarity, and FPRA.

### Benchmarking of phage profiling using CAMI *in silico* datasets

Phage profiling has emerged as an underexplored domain, crucial for delineating bacteria-phage interactions within the microbiome. The field has few extensively validated tools for accurate phage detection. To address this gap, the capabilities of CHAMP for phage profiling were rigorously evaluated against the current state-of-the-art phage profiler, Phanta (Pinto et al., 2023), which, like CHAMP, leverages k-mer matching for virus identification. The benchmarking dataset (n = 10) was simulated to reflect a realistic composition of 95% bacteria and 5% bacteriophages, with sequencing depths between 10 and 50 million reads, corresponding to the benchmarking framework previously proposed by Pinto et al., 2023. To ensure a balanced comparison, only genomes with species-level annotations shared between the databases were included. Phanta showed good precision (avg. 0.95 ± 0.02); however it was surpassed by CHAMP, which demonstrated near-perfect precision (avg. 0.99 ± 0.01, Figure 4A). CHAMP also outperformed Phanta in recall (avg. 0.95 ± 0.02 vs. 0.84 ± 0.05), F1 scores (avg. 0.97 ± 0.01 vs. 0.87 ± 0.03), and similarity (0.98 ± 0.01 vs. 0.90 ± 0.07), while maintaining a lower FPRA (avg. 0.07% ± 0.13% vs. 5.08% ± 3.34%, Figure 4B). CHAMP therefore exhibited superior performance across all benchmarking metrics, demonstrating both high precision and substantial recall, thereby establishing its precedence in phage profiling accuracy.

**Figure 4.**
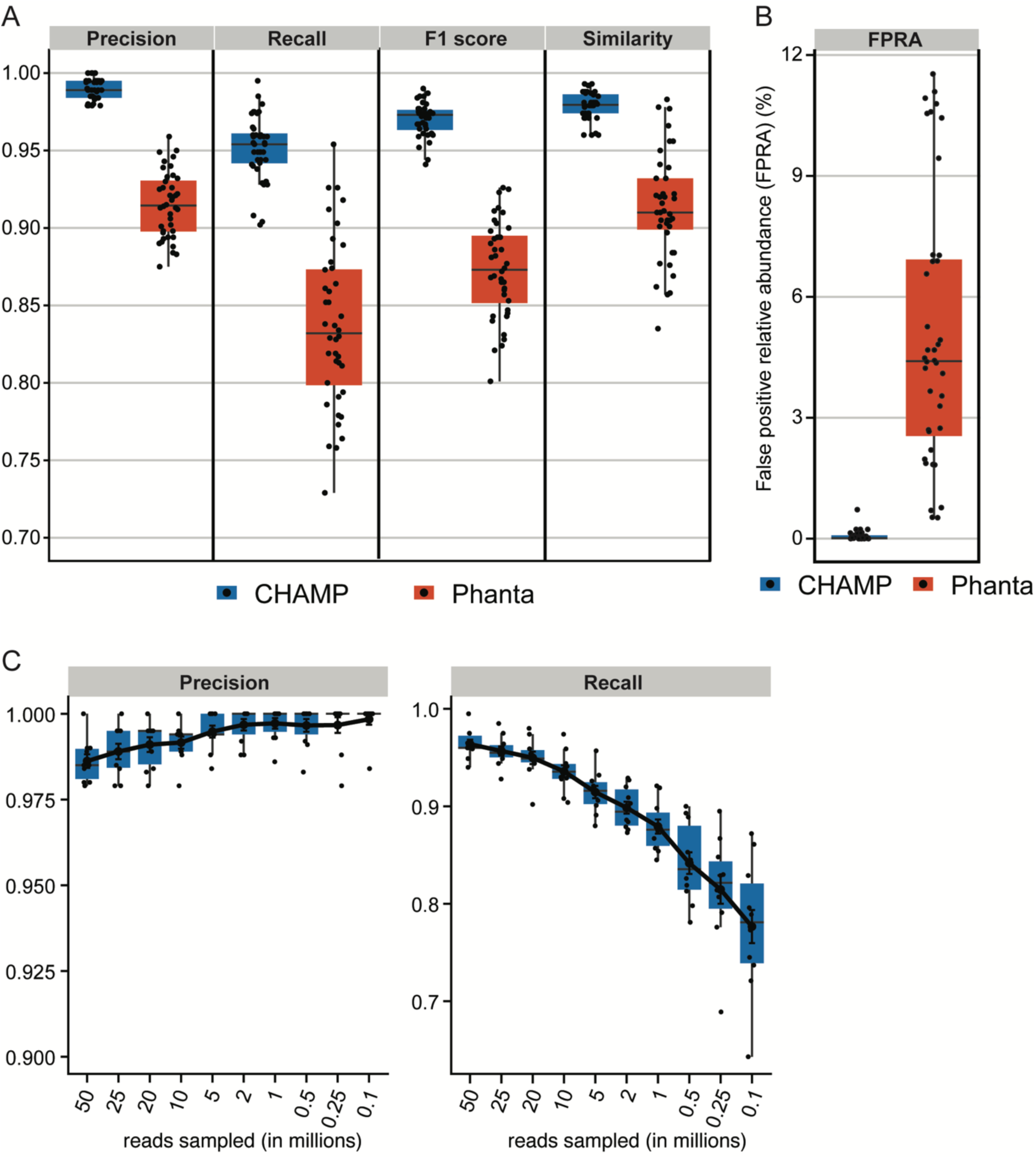
(**A-B)** Benchmark of phage profiling on *in silico* dataset with 10 simulated metagenomes containing 95% prokaryotic and 5% phage reads across the sequencing depths: 10, 20, 25, and 50 million reads. The performance of phage profiling with CHAMP and Phanta was assessed by F1 score, precision, recall, similarity, and FPRA. (**C)** Precision and recall of CHAMP after rarefying the 10 *in silico* communities with 50 million to 25, 20, 10, 5, 2, 1, 0.5, 0.25, and 0.1 million reads.

Phanta does not recommend phage profiling in samples below 10 million reads. Nonetheless, CHAMP maintained exceptional precision (>0.98) across a broad range of sequencing depths, of which the lowest depth had only 100,000 reads (Figure 4C). Recall rates decreased with reduced sequencing depth, mirroring observations in prokaryotic community benchmarks. Recall remained high (>0.75) even at a depth of only 100,000 reads, underscoring the superior performance of CHAMP in profiling phage communities across deep and shallow samples.

## Discussion

CHAMP was developed to accurately profile all types of microbes including eukaryotes, archaea, bacteria, and viruses found in human microbiome samples. Despite the existence of multi-domain and phage-inclusive profilers (Parks et al., 2021; Zolfo et al., 2024), CHAMP is the first to systematically evaluate the performance across prokaryotes, eukaryotes, and phages, thereby providing a broad evaluation and comparison of current profilers. When compared against MetaPhlAn 4, mOTUs 3, Bracken 2, and Phanta, CHAMP achieved the highest recall and F1 scores of all tools across all benchmark datasets (Figure 1, 3, and 4). CHAMP also consistently achieved the lowest FPRA that was up to 110 times lower than the second-best tool, emphasizing its superior accuracy in the taxonomic profiling of diverse and complex microbiome samples.Accounting for differences in database comprehensiveness is crucial to ensure fairness in the comparison of profiling tools. The prokaryote profiling accuracy of CHAMP was assessed against MetaPhlAn 4, mOTUs 3, and Bracken 2, and based on two reference datasets from NIBSC with common gut microbiome species and 50 *in silico* metagenomes. The *in silico* metagenomes were designed to reflect the diversity across different body sites, including human adult gut, infant gut, oral, and skin samples. The species included in each body sites were selected from relevant publications to ensure fairness in the benchmark. For the eukaryotic and phage benchmarks, where CHAMP was compared against MetaPhlAn 4 and Phanta, respectively, the communities were simulated using the subset of species or genome sequences shared between the databases. As CHAMP was developed specifically for profiling the human microbiome, it should not be used to profile metagenomes from non-human origin.

A disadvantage to limiting the species in the benchmark to cultured species shared across databases is that unknown members of the microbiome are not included. Other benchmarks have focussed on covering primarily unknown species (Parks et al., 2021). CHAMP was built with the intention of profiling rare and/or uncultured species as evident from our large efforts in collecting and building MAGs from 15,224 samples from under-sampled body sites and countries. However, we did not include unknown species for two reasons: first, to make the benchmark fair across different tools and databases as some databases may not include uncultured species and second, because we designed body-site specific communities using published relative abundance matrices where only cultured species were profiled.

Accounting for differences in species present in different databases does not mean the comprehensiveness and quality of the database does not affect profiling accuracy. For example, Bracken 2 showed promising results on DNA reference reagents, presumably because the reagents consisted of well-characterized species. Nonetheless, Bracken 2 underperformed on the *in-silico* communities using its suggested database, RefSeq (2023-09-10). Still, the performance of Bracken 2 did not improve when we sought to create a more comprehensive k-mer database based on GTDB (Supplementary Figure 1). Bracken 2 using GTDB as a database also resulted in lower sensitivity on the DNA reagents, Gut-Hi-Low-RR and Gut-Mix-RR, which was 35 for both, compared to 100 and 91.6 for Bracken 2 built with the standard RefSeq database (data not shown). Some authors have noted parameters used and database choice in Bracken 2 influences predicted profiles (Wright et al., 2023). Although we tinkered with the Bracken 2 database for benchmarking purposes, optimizing Bracken 2 run parameters would be unfair to the other profilers evaluated as they were all run using recommended settings.

Runtime and memory usage often affect the choice of profilers used (Meyer et al., 2022). In this regard, k-mer based methods (e.g. Bracken 2) are typically faster but use more memory compared to alignment-dependent methods, such as CHAMP, as can be seen in Supplementary Tables 5-6.

As CHAMP uses a different profiling methodology for bacteriophage profiling, we also evaluated bacteriophage profiling resource use compared to Phanta on a single viral metagenome. CHAMP bacteriophage profiling was also significantly slower than Phanta (2 hrs 16 mins vs 36 mins 40 secs) but displayed significantly less peak memory usage (0.09 Gb vs 33.27 Gb). CHAMP was designed for parallelization, so just enabling profiling on 4 threads would bring total runtime under an hour, whilst bringing total maximum memory to around 14Gb. In this scenario, its runtime and maximum memory usage becomes very similar to MetaPhlAn 4. All in all, the time and memory available needed to run CHAMP does not significantly differ from the other considered profilers, as there is a clear tradeoff between runtime and maximum memory use. CHAMP can be run on the Cosmos-ID hub (https://cosmosidhub.com/) and by default, takes about an hour to process a sample of 20 million reads. 10 samples can be run in parallel.

CHAMP is solely intended for profiling the human microbiome, whilst the microbiome profilers we benchmarked against are designed to profile human, host-associated, and environmental microbiomes. As such, the background noise of these profilers when performing human microbiome profiling will be higher than CHAMP, as they must distinguish among more species increasing the likelihood of species misclassification. Another key to CHAMP performance is the number of human specific references and the curation of the reference database. The database was constructed using strict quality filtering of MAGs and (completeness >= 90%, contamination < 0.05 and multiple tests for chimerism, for details see Materials and Methods), to improve the quality of the signature genes (Orakov et al., 2021). MetaPhlAn 4 and mOTUs 3 include assemblies with completeness values >= 50% and contamination values < 5% and 10% into their databases, respectively. These differences likely explain the consistently higher recall and lower FPRA that CHAMP displays across benchmarks. Moreover, CHAMP employs a dynamic sample-specific signal filtering (for details see Materials and Methods) that handles spurious mapping, further reduces false signal, and improves precision and abundance estimates.

Signature or marker gene-based profiling strategies, such as those employed by MetaPhlAn 4 and CHAMP for prokaryotic and eukaryotic profiling come with its challenges. At times, signature genes may be non-specific to a species or absent in a given sample. Hence, CHAMP implements a dynamic sample-specific signature gene filtering approach to reduce FPs as well as improve abundance estimates. CHAMP maps all reads of a given sample to the entire HMR05 gene catalogue (pan genomes) to obtain the total species read count. Then, it models the expected read counts mapping to signature genes using a negative binomial distribution. The expectation is that as the number of reads for that species increases, the number of reads mapped to its signature genes also increases until all signature genes are captured. Signature genes outside the 99% quantile distribution representing non-specific genes at the top or missing genes at the bottom of the distribution are removed. We believe the thorough profiling strategy of CHAMP explains its lower FPRA estimates compared to the next-best profiler MetaPhlAn 4, which adopts a more rigid cutoff-based approach. It estimates abundance by removing the upper and lower 20% of marker gene coverage estimates to tackle non-specific marker genes and the random absence of marker genes, respectively. The quality of the HMR05 database coupled with its unique signature gene approach may help explain why CHAMP sees the lowest impact of FPs across profilers while still achieving high precision and similarity scores.

Overall, CHAMP performed better than other tools in terms of precision and similarity, and in specific benchmarks at least at par with other profilers. For the prokaryotic and eukaryotic *in silico* benchmarks, CHAMP and MetaPhlan 4 both performed extremely well with CHAMP outperforming MetaPhlAn 4 across 3 out of 5 body sites for prokaryotic profiling as well as for eukaryotic profiling. Likewise, CHAMP and MetaPhlAn 4 had the highest similarity values for the *in silico* prokaryotic communities as well as comparable similarity scores on the DNA reference reagents, where Bracken 2 achieved marginally better scores. Eukaryotic and viral profiling are less mature than prokaryotic profiling with a lower number of methodologies published. The fact that CHAMP consistently outperformed MetaPhlAn 4 and Phanta across all benchmarking metrics when it came to profiling eukaryotes (MetaPhlAn 4) and viruses (Phanta) shows that CHAMP paves the way for a new generation of taxonomic profilers that are also able to capture viral and eukaryotic members of the microbiome.

The exploration of sensitivity thresholds in taxonomic profiling rarely occurs in benchmarking studies, yet it is crucial, especially as profiling accuracy is lower for low abundant species or samples with low microbial biomass. An example of such is the suboptimal profiling of tumor biopsies where inaccurate detection of species led to the conclusion that different cancer types had their own unique microbiome (Gihawi et al., 2023). We explored the sensitivity of CHAMP along with MetaPhlan 4, mOTUs 3, and Bracken 2 by comparing their accuracy to diminishing sequencing depths. CHAMP proved to be more robust at lower sequencing depths than all other profilers (Figures 2 and 4C), with very high recall and low false discovery rates even at very shallow sequencing depths. Taken together, benchmarks show that CHAMP sets a new standard for the accuracy of multi-domain and phage-inclusive human microbiome profiling across sample types and body sites. CHAMP software and code is proprietary, but it is now available online for a fee; https://cosmosidhub.com.

## Author Contributions

All authors took part in the development of CHAMP. SP, JJ, and JR performed the benchmarking. SP, JJ, and PNM prepared the first draft of the manuscript. All authors took part in the editing of the manuscript.

## Acknowledgments

We would like to thank several people for contributing to the discussion of the development of CHAMP, including Trine Zachariasen, Gisle Vestergaard, Myeonghyun Yoou, Dennis Pohl, Albert Pallejà, Oksana Lukjancenko, Janne Marie Moll, and Thorsten Brach. Thanks to Abril Diosdado Hernandez for bioinformatics support.

## Data Availability Statement

Sequencing data and ground truth taxonomic profiles for *in silico* mock communities generated have been deposited to Zenodo: 10.5281/zenodo.10777404 (prokaryotes and phages) and 10.5281/zenodo.12090449 (eukaryotes). Sequencing data for NIBSC DNA mock communities, Gut-Mix-RR and Gut-HiLo-RR can be found under BioProject ID PRJNA622674.

---

**Supplementary Figure 1.**
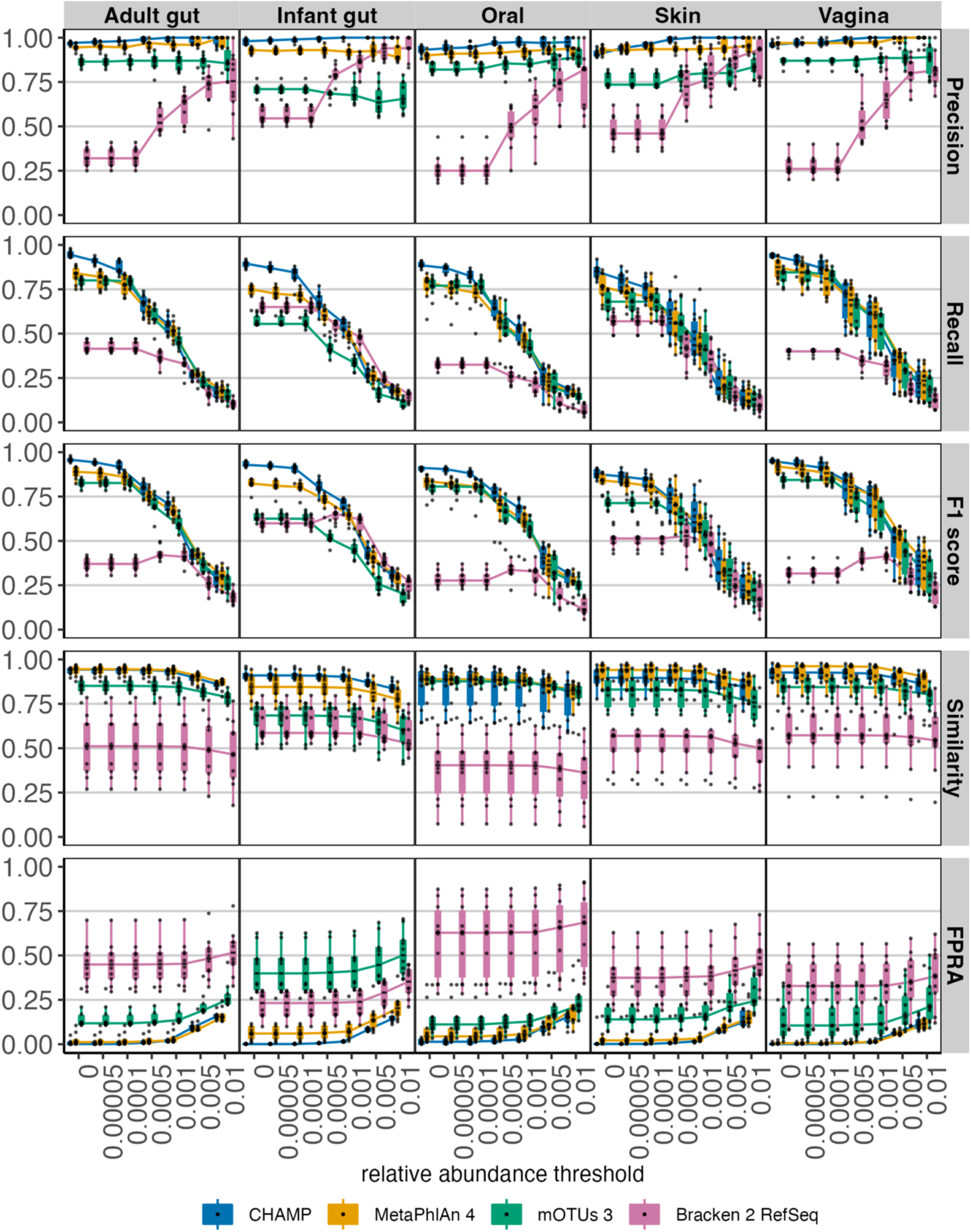
Detection limits at species relative abundance cutoffs of 0.0005, 0.0001, 0.0005, 0.001, 0.005 and 0.001. 50 metagenomes were used and subset at the abundance cutoffs leading to 350 communities.

**Supplementary Figure 2.**
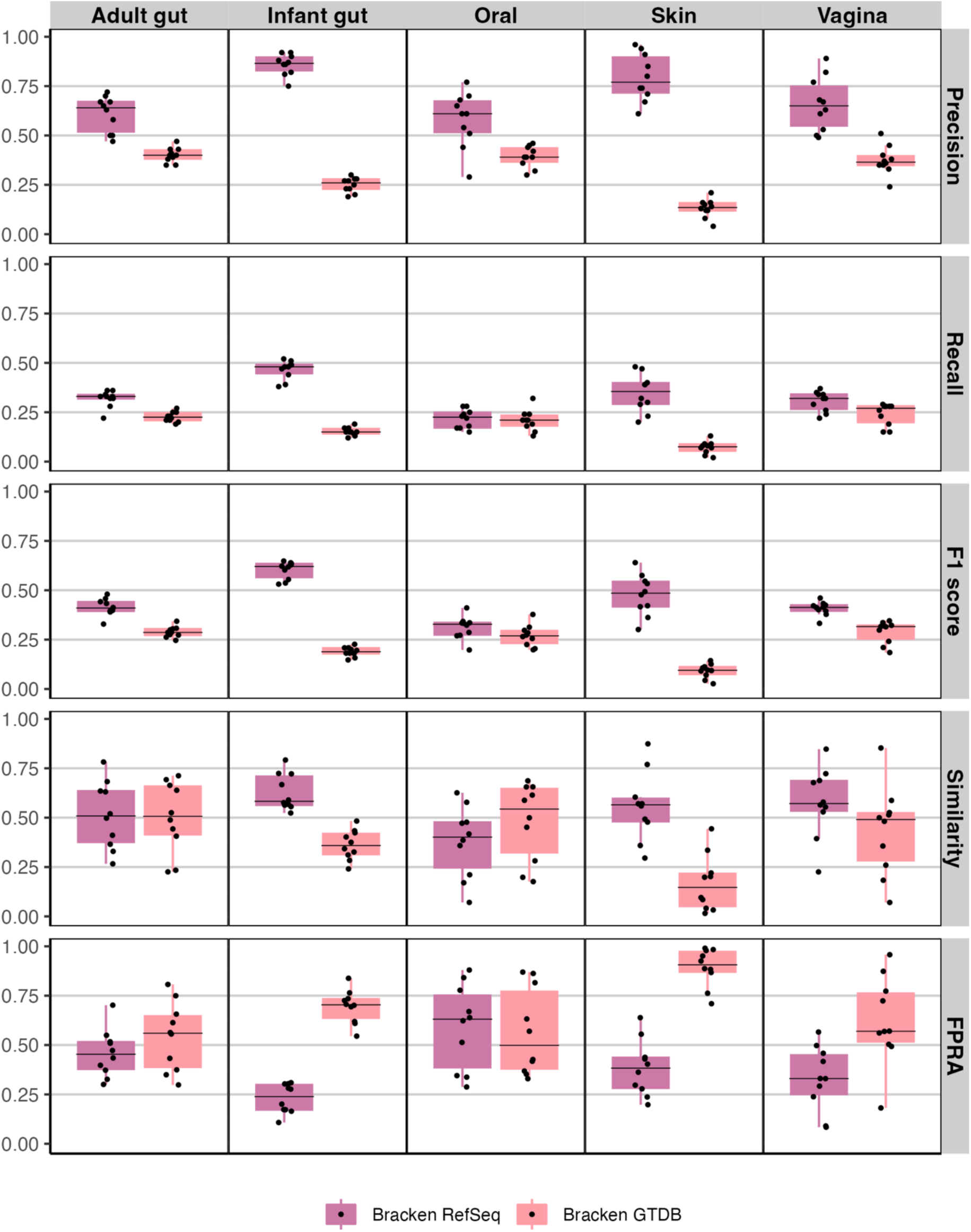
Evaluating Bracken 2 with RefSeq vs GTDB r214 on key benchmarking metrics across 5 human body communities. 50 metagenomic communities (10 per body site) comprising 100 prokaryotic species each were used.

**Supplementary Figure 3.**
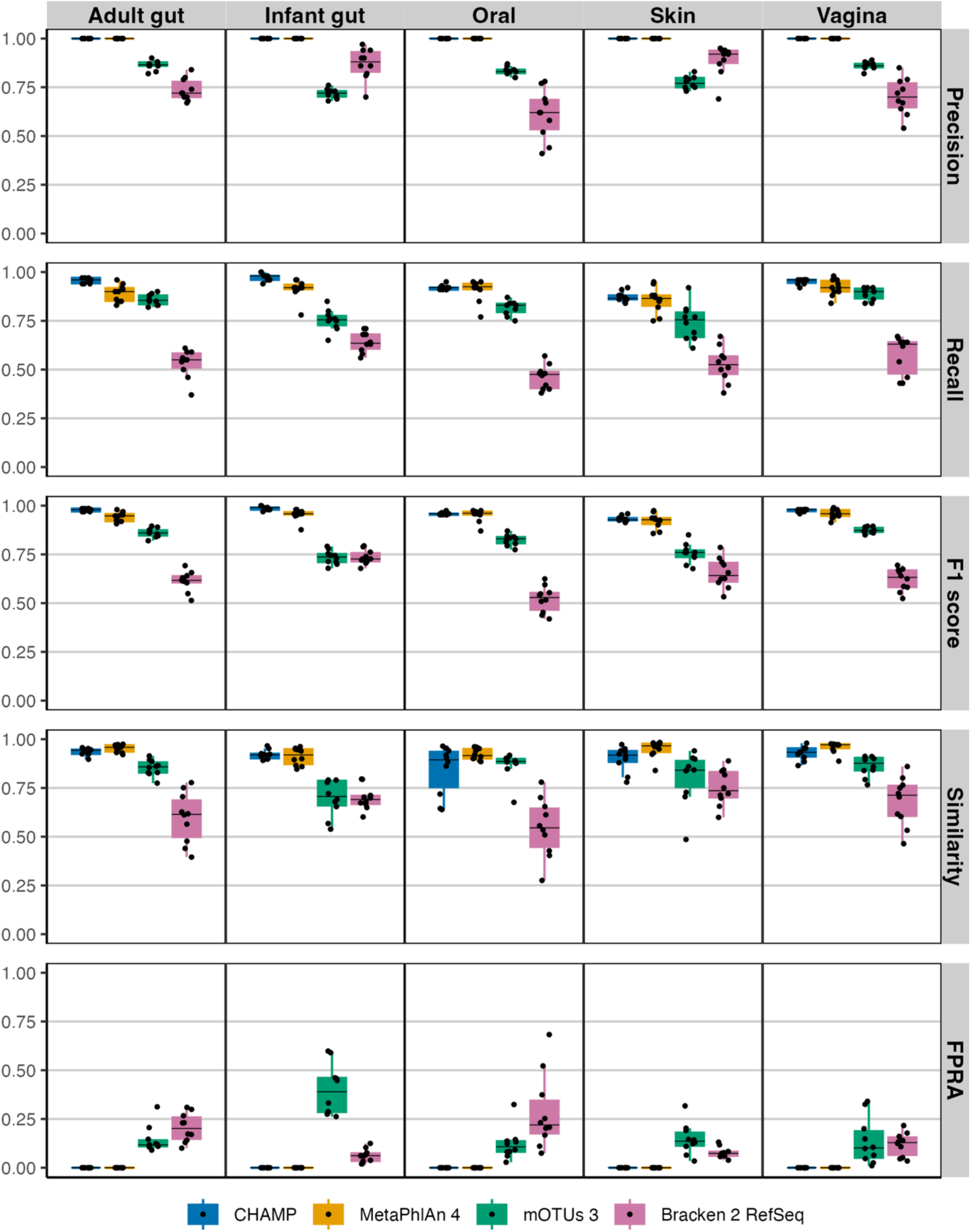
Genus-level **e**valuation the taxonomic profilers CHAMP, MetaPhlAn 4, mOTUs 3, and Bracken 2 on key benchmarking metrics: precision, recall, F1 score, similarity, and FPRA across five human body communities. A total of 50 metagenomes (10 per body site) each with 100 prokaryotic species were simulated using CAMISIM.

## Supplementary Tables

**Supplementary Table 1.** Genome accessions used for *in silico* metagenome simulations across body site communities

**Supplementary Table 2.** Genome accessions used for eukaryotes *in silico* metagenome simulations

**Supplementary Table 3.** Genome accessions used for virome *in silico* metagenome simulations

**Supplementary Table 4.** Overview of benchmarking metrics for *in silico* body site metagenome communities across profilers

**Supplementary Table 5.** Runtime and maximum memory comparisons (in Gb) comparisons across profilers for one *in silico* gut adult metagenome community (2.1 Gb). All profilers were run on default settings in a 1 compute node (40 processors per node) HPC.

**Supplementary Table 6.** Runtime and maximum memory comparisons (in Gb) comparisons between CHAMP and Phanta for bacteriophage profiling using one 50M *in silico* metagenome. All profilers were run on default settings in a 1 compute node (40 processors per node) HPC.

## References

Almeida, A., Mitchell, A. L., Boland, M., Forster, S. C., Gloor, G. B., Tarkowska, A., et al. (2019). A new genomic blueprint of the human gut microbiota. Nature 568, 499–504. doi: 10.1038/S41586-019-0965-1

Almeida, A., Nayfach, S., Boland, M., Strozzi, F., Beracochea, M., Shi, Z. J., et al. (2020). A unified catalog of 204,938 reference genomes from the human gut microbiome. Nature Biotechnology 2020 39:1 39, 105–114. doi: 10.1038/s41587-020-0603-3

Amos, G. C. A., Logan, A., Anwar, S., Fritzsche, M., Mate, R., Bleazard, T., et al. (2020). Developing standards for the microbiome field. Microbiome 8, 1–13. doi: 10.1186/S40168-020-00856-3/FIGURES/3

Bäckhed, F., Roswall, J., Peng, Y., Feng, Q., Jia, H., Kovatcheva-Datchary, P., et al. (2015). Dynamics and Stabilization of the Human Gut Microbiome during the First Year of Life. Cell Host Microbe 17, 690–703. doi: 10.1016/J.CHOM.2015.04.004

Benler, S., Yutin, N., Antipov, D., Rayko, M., Shmakov, S., Gussow, A. B., et al. (2021). Thousands of previously unknown phages discovered in whole-community human gut metagenomes. Microbiome 2021 9:1 9, 1–17. doi: 10.1186/S40168-021-01017-W

Blanco-Míguez, A., Beghini, F., Cumbo, F., McIver, L. J., Thompson, K. N., Zolfo, M., et al. (2023). Extending and improving metagenomic taxonomic profiling with uncharacterized species using MetaPhlAn 4. Nature Biotechnology 2023 41:11 41, 1633–1644. doi: 10.1038/s41587-023-01688-w

Camarillo-Guerrero, L. F., Almeida, A., Rangel-Pineros, G., Finn, R. D., and Lawley, T. D. (2021). Massive expansion of human gut bacteriophage diversity. Cell 184, 1098. doi: 10.1016/J.CELL.2021.01.029

Dekkers, K. F., Sayols-Baixeras, S., Baldanzi, G., Nowak, C., Hammar, U., Nguyen, D., et al. (2022). An online atlas of human plasma metabolite signatures of gut microbiome composition. Nature Communications 2022 13:1 13, 1–12. doi: 10.1038/s41467-022-33050-0

Fritz, A., Hofmann, P., Majda, S., Dahms, E., Dröge, J., Fiedler, J., et al. (2019). CAMISIM: Simulating metagenomes and microbial communities. Microbiome 7, 1–12. doi: 10.1186/S40168-019-0633-6/FIGURES/5

Gihawi, A., Ge, Y., Lu, J., Puiu, D., Xu, A., Cooper, C. S., et al. (2023). Major data analysis errors invalidate cancer microbiome findings. mBio 14. doi: 10.1128/MBIO.01607-23

Huttenhower, C., Gevers, D., Knight, R., Abubucker, S., Badger, J. H., Chinwalla, A. T., et al. (2012). Structure, function and diversity of the healthy human microbiome. Nature 2012 486:7402 486, 207–214. doi: 10.1038/nature11234

Johansen, J., Plichta, D. R., Nissen, J. N., Jespersen, M. L., Shah, S. A., Deng, L., et al. (2022). Genome binning of viral entities from bulk metagenomics data. Nat Commun 13. doi: 10.1038/S41467-022-28581-5

Liang, G., and Bushman, F. D. (2021). The human virome: assembly, composition and host interactions. Nature Reviews Microbiology 2021 19:8 19, 514–527. doi: 10.1038/s41579-021-00536-5

Lu, J., Breitwieser, F. P., Thielen, P., and Salzberg, S. L. (2017). Bracken: Estimating species abundance in metagenomics data. PeerJ Comput Sci 2017, e104. doi: 10.7717/PEERJ-CS.104/SUPP-5

Meyer, F., Bremges, A., Belmann, P., Janssen, S., McHardy, A. C., and Koslicki, D. (2019). Assessing taxonomic metagenome profilers with OPAL. Genome Biol 20, 1–10. doi: 10.1186/S13059-019-1646-Y/FIGURES/3

Meyer, F., Fritz, A., Deng, Z. L., Koslicki, D., Lesker, T. R., Gurevich, A., et al. (2022). Critical Assessment of Metagenome Interpretation: the second round of challenges. Nature Methods 2022 19:4 19, 429–440. doi: 10.1038/s41592-022-01431-4

Nayfach, S., Shi, Z. J., Seshadri, R., Pollard, K. S., and Kyrpides, N. C. (2019). New insights from uncultivated genomes of the global human gut microbiome. Nature 568, 505–510. doi: 10.1038/S41586-019-1058-X

Nissen, J. N., Johansen, J., Allesøe, R. L., Sønderby, C. K., Armenteros, J. J. A., Grønbech, C. H., et al. (2021). Improved metagenome binning and assembly using deep variational autoencoders. Nat Biotechnol 39, 555–560. doi: 10.1038/s41587-020-00777-4

O’Leary, N. A., Wright, M. W., Brister, J. R., Ciufo, S., Haddad, D., McVeigh, R., et al. (2016). Reference sequence (RefSeq) database at NCBI: current status, taxonomic expansion, and functional annotation. Nucleic Acids Res 44, D733–D745. doi: 10.1093/NAR/GKV1189

Olm, M. R., West, P. T., Brooks, B., Firek, B. A., Baker, R., Morowitz, M. J., et al. (2019). Genome-resolved metagenomics of eukaryotic populations during early colonization of premature infants and in hospital rooms. Microbiome 7, 1–16. doi: 10.1186/S40168-019-0638-1/FIGURES/4

Orakov, A., Fullam, A., Coelho, L. P., Khedkar, S., Szklarczyk, D., Mende, D. R., et al. (2021). GUNC: detection of chimerism and contamination in prokaryotic genomes. Genome Biol 22. doi: 10.1186/S13059-021-02393-0

Parks, D. H., Rigato, F., Vera-Wolf, P., Krause, L., Hugenholtz, P., Tyson, G. W., et al. (2021). Evaluation of the Microba Community Profiler for Taxonomic Profiling of Metagenomic Datasets From the Human Gut Microbiome. Front Microbiol 12, 731. doi: 10.3389/FMICB.2021.643682/BIBTEX

Pasolli, E., Asnicar, F., Manara, S., Zolfo, M., Karcher, N., Armanini, F., et al. (2019). Extensive Unexplored Human Microbiome Diversity Revealed by Over 150,000 Genomes from Metagenomes Spanning Age, Geography, and Lifestyle. Cell 176, 649–662.e20. doi: 10.1016/J.CELL.2019.01.001

Pinto, Y., Chakraborty, M., Jain, N., and Bhatt, A. S. (2023). Phage-inclusive profiling of human gut microbiomes with Phanta. Nat Biotechnol. doi: 10.1038/S41587-023-01799-4

Poussin, C., Khachatryan, L., Sierro, N., Narsapuram, V. K., Meyer, F., Kaikala, V., et al. (2022). Crowdsourced benchmarking of taxonomic metagenome profilers: lessons learned from the sbv IMPROVER Microbiomics challenge. BMC Genomics 23, 624. doi: 10.1186/S12864-022-08803-2/FIGURES/7

Proctor, L. (2019). Priorities for the next 10 years of human microbiome research. Nature 2021 569:7758 569, 623–625. doi: 10.1038/d41586-019-01654-0

Ruscheweyh, H. J., Milanese, A., Paoli, L., Karcher, N., Clayssen, Q., Keller, M. I., et al. (2022). Cultivation-independent genomes greatly expand taxonomic-profiling capabilities of mOTUs across various environments. Microbiome 10, 1–12. doi: 10.1186/S40168-022-01410-Z/FIGURES/5

Saheb Kashaf, S., Proctor, D. M., Deming, C., Saary, P., Hölzer, M., Mullikin, J., et al. (2021). Integrating cultivation and metagenomics for a multi-kingdom view of skin microbiome diversity and functions. Nature Microbiology 2021 7:1 7, 169–179. doi: 10.1038/s41564-021-01011-w

Saraiva, J. P., Bartholomäus, A., Toscan, R. B., Baldrian, P., and Nunes da Rocha, U. (2023). Recovery of 197 eukaryotic bins reveals major challenges for eukaryote genome reconstruction from terrestrial metagenomes. Mol Ecol Resour 23, 1066–1076. doi: 10.1111/1755-0998.13776

Schoch, C. L., Ciufo, S., Domrachev, M., Hotton, C. L., Kannan, S., Khovanskaya, R., et al. (2020). NCBI Taxonomy: a comprehensive update on curation, resources and tools. Database (Oxford) 2020. doi: 10.1093/DATABASE/BAAA062

Sczyrba, A., Hofmann, P., Belmann, P., Koslicki, D., Janssen, S., Dröge, J., et al. (2017). Critical Assessment of Metagenome Interpretation—a benchmark of metagenomics software. Nature Methods 2017 14:11 14, 1063–1071. doi: 10.1038/nmeth.4458

Shah, S. A., Deng, L., Thorsen, J., Pedersen, A. G., Dion, M. B., Castro-Mejía, J. L., et al. (2023). Expanding known viral diversity in the healthy infant gut. Nature Microbiology 2023 8:5 8, 986–998. doi: 10.1038/s41564-023-01345-7

Sun, Z., Huang, S., Zhang, M., Zhu, Q., Haiminen, N., Carrieri, A. P., et al. (2021). Challenges in benchmarking metagenomic profilers. Nature Methods 2021 18:6 18, 618–626. doi: 10.1038/s41592-021-01141-3

Wood, D. E., Lu, J., and Langmead, B. (2019). Improved metagenomic analysis with Kraken 2. Genome Biol 20, 1–13. doi: 10.1186/S13059-019-1891-0/FIGURES/2

Wright, R. J., Comeau, A. M., and Langille, M. G. I. (2023). From defaults to databases: parameter and database choice dramatically impact the performance of metagenomic taxonomic classification tools. Microb Genom 9, 000949. doi: 10.1099/MGEN.0.000949/CITE/REFWORKS

Zeng, S., Patangia, D., Almeida, A., Zhou, Z., Mu, D., Paul Ross, R., et al. (2022). A compendium of 32,277 metagenome-assembled genomes and over 80 million genes from the early-life human gut microbiome. Nature Communications 2022 13:1 13, 1–15. doi: 10.1038/s41467-022-32805-z

Zmora, N., Zilberman-Schapira, G., Suez, J., Mor, U., Dori-Bachash, M., Bashiardes, S., et al. (2018). Personalized Gut Mucosal Colonization Resistance to Empiric Probiotics Is Associated with Unique Host and Microbiome Features. Cell 174, 1388–1405.e21. doi: 10.1016/J.CELL.2018.08.041

Zolfo, M., Silverj, A., Blanco-Míguez, A., Manghi, P., Rota-Stabelli, O., Heidrich, V., et al. (2024). Discovering and exploring the hidden diversity of human gut viruses using highly enriched virome samples. bioRxiv, 2024.02.19.580813. doi: 10.1101/2024.02.19.580813

